# Depolarization induces calcium-dependent BMP4 release from mouse embryonic palate mesenchyme

**DOI:** 10.1101/2024.06.11.598333

**Authors:** Mikaela L Follmer, Trevor Isner, Yunus H. Ozekin, Claire Levitt, Emily Anne Bates

**Author notes:** **Corresponding Author:** Emily Anne Bates, University of Colorado Anschutz Medical Campus, 12800 E 19^th^ Avenue, Aurora, CO 80045. indicates equal contribution (Mikaela L Follmer, Trevor Isner, Yunus H. Ozekin).

## Abstract

Ion channels are essential for proper morphogenesis of the craniofacial skeleton. However, the molecular mechanisms underlying this phenomenon are unknown. Loss of the *Kcnj2* potassium channel disrupts Bone Morphogenetic Protein (BMP) signaling within the developing palate. BMP signaling is essential for the correct development of several skeletal structures, including the palate, though little is known about the mechanisms that govern BMP secretion. We introduce a tool to image the release of bone morphogenetic protein 4 (BMP4) from mammalian cells. Using this tool, we show that depolarization induces BMP4 release from mouse embryonic palate mesenchyme cells in a calcium-dependent manner. We show native transient changes in intracellular calcium occur in cranial neural crest cells, the cells from which embryonic palate mesenchyme derives. Waves of transient changes in intracellular calcium suggest that these cells are electrically coupled and may temporally coordinate BMP release. These transient changes in intracellular calcium persist in palate mesenchyme cells from embryonic day (E) 9.5 to 13.5 mice. Disruption of *Kcnj2* significantly decreases the amplitude of calcium transients and the ability of cells to secrete BMP. Together, these data suggest that temporal control of developmental cues is regulated by ion channels, depolarization, and changes in intracellular calcium for mammalian craniofacial morphogenesis.

**SUMMARY:** We show that embryonic palate mesenchyme cells undergo transient changes in intracellular calcium. Depolarization of these cells induces BMP4 release suggesting that ion channels are a node in BMP4 signaling.

## INTRODUCTION

Genetic syndromes demonstrate that ion channels contribute to human facial morphogenesis. Syndromes caused by disruption or activation of ion channel function are termed channelopathies. An analysis of these disorders demonstrates abnormal craniofacial development is associated with mutations that disrupt calcium channels (e.g. *CACNA1C*, Timothy syndrome) (*1–3*), potassium channels (e.g. *KCNK9*-Birk-Barel syndrome; *KCNJ2*-Andersen-Tawil syndrome, *GIRK2*-Keppen-Ludinski syndrome) (*4–14*), and sodium channels (e.g. *NALCN*-Infantile hypotonia with psychomotor retardation and characteristic faces (IHPRF))(*15*). Furthermore, intrauterine exposure to teratogens that impact ion channel function, such as anti-epileptic drugs, heat, nicotine, and cannabinoids, is associated with increased risk for craniofacial defects(*15–25*). Similarly, genetic inhibition of ion channels can cause morphological abnormalities in several species of animals (*26–33*). Ion channels work together to establish the electrical properties, including membrane potential of each cell. The diversity of ion channels that are important for craniofacial development suggests that cellular electrical properties impact morphogenesis. However, the molecular mechanisms by which membrane potential contributes to craniofacial morphogenesis remain unclear.

One of several possibilities is that ion channels impact the complex communication patterns between cells during facial morphogenesis. Proper formation of the face requires cranial neural crest (CNC) cells to migrate from the neural tube to populate the frontal nasal process and pharyngeal arches (*34, 35*). CNC cells form the craniofacial bone and cartilage, cranial neurons and glia, odontoblasts, and melanocytes and require precise spatiotemporal control to produce the proper cell types (*36*). This is accomplished through signaling networks of morphogens, such as Bone morphogenetic protein (BMP), Notch, and Sonic hedgehog (Shh) (*37, 38*). An impressive amount of research has revealed the underpinnings of these molecular signaling cascades. For example, BMP is an essential molecular signal for craniofacial development. Upon BMP ligand binding, receptors phosphorylate Smads which can then enter the nucleus to induce target gene expression. BMP signaling also activates extracellular signal-regulated kinases (ERKs) for osteoblast differentiation (*39*). Oscillatory, or pulsatile ligand exposure is key for BMP to communicate efficiently via receptor engagement and Smad activation (*40*). Osteoblast development during regeneration relies on waves of oscillatory ERK activation in Zebrafish (*41*) suggesting that temporal regulation is important for both modes of BMP signaling. However, we know very little about how cells control the timing of molecular signals that mediate BMP release. If the temporal signaling pattern matters for the transcriptional output, how do cells control ligand release? One possibility is that ion channels control pulsatile cellular release of ligands.

The established roles of ion channels in traditionally excitable cell types, such as neurons, may lend insight into the role of membrane potential in craniofacial development. In neurons, ion channels coordinate the precise release of vesicles containing molecular signals, called neurotransmitters, to orchestrate complex communication between cells (*42*). This is accomplished when the cell’s membrane potential rapidly changes due to the influx of sodium and calcium, and the efflux of potassium ions controlled by the opening and closing of ion channels. Thus, an electrical signal coordinates the delivery of a chemical signal. Recent evidence suggests that ion channels may perform a similar function to control timing and release of developmental cues in cell types not classically thought of as excitable. Inwardly rectifying potassium channel *KCNJ2* (Kir2.1) loss of function in humans, mice, and flies causes defects that are remarkably similar to those arising from a loss of BMP signaling (*27, 29*). In *Drosophila*, a homolog of *Kcnj2* called Irk2 is required for the BMP homolog Decapentaplegic (Dpp)-mediated patterning of the wing (*29*). Loss of Irk2 conduction disrupts pulsatile Dpp release in the wing primordium (wing disc) (*28*). Further, Dpp release can be induced by depolarization (*28*). In cell culture, pulsatile presentation of BMP ligands produces a greater transcriptional response than constant exposure (*40*), suggesting an intimate link between coordinated timing of release and downstream patterning. This requirement for Kcnj2 homologs in BMP signaling is conserved, as *Kcnj2* is important for efficient BMP signaling in mice (*27*). *Kcnj2* and BMP are required in cranial neural crest cells for craniofacial patterning (*27, 43*). These data suggest the enticing possibility that membrane potential controls BMP release with a similar mechanism to neuronal neurotransmitter release. Ion channels regulate membrane potential, which controls calcium-dependent release of vesicles containing developmental signals.

Here, we investigate the hypothesis that ion channel-mediated membrane potential controls BMP release from palatal mesenchyme cells. We introduce a novel tool to visualize BMP4 release from cells from the developing mouse palatal mesenchyme. We test if depolarization of mouse embryonic palatal mesenchyme cells induces vesicular fusion of BMP-containing vesicles and if this process is calcium-dependent. We show that depolarization stimulates BMP4 release from these cells. We demonstrate that CNC cells and embryonic palate mesenchyme cells exhibit endogenous transient changes in intracellular calcium that can propagate through waves between cells. Calcium transients are dependent on Kcnj2. Together our results suggest ion channels can act upstream of BMP signaling in the developing mouse palate.

## RESULTS

### Membrane potential regulates BMP release

Mutations in Kcnj2 cause cleft palate and other craniofacial defects and reduce BMP signaling in multiple organisms (9, 26–29, 44). Because Kcnj2 regulates resting membrane potential, we use it as a tool to determine how membrane potential contributes to BMP signaling. We hypothesized that Kcnj2 regulates BMP ligand release. To test this hypothesis, we incubated WT mouse embryonic fibroblasts in conditioned media from WT, *Kcnj2^ko/+^*, or *Kcnj2^koko^* cultured mouse embryonic fibroblasts (Figure 1A). We then measured phosphorylated Smad 1/5 in the receiving cells as a readout of BMP ligands in the conditioned media (Figure 1A). Smad 1/5 phosphorylation was significantly reduced in cells treated with *Kcnj2^ko/ko^* and *Kcnj2^ko/+^* conditioned media compared to cells treated with WT-conditioned media (Figure 1B-C). These data are consistent with the hypothesis that membrane potential regulates BMP ligand release. However, we needed a method to measure BMP release from cells to further investigate our hypothesis.

**Figure 1:**
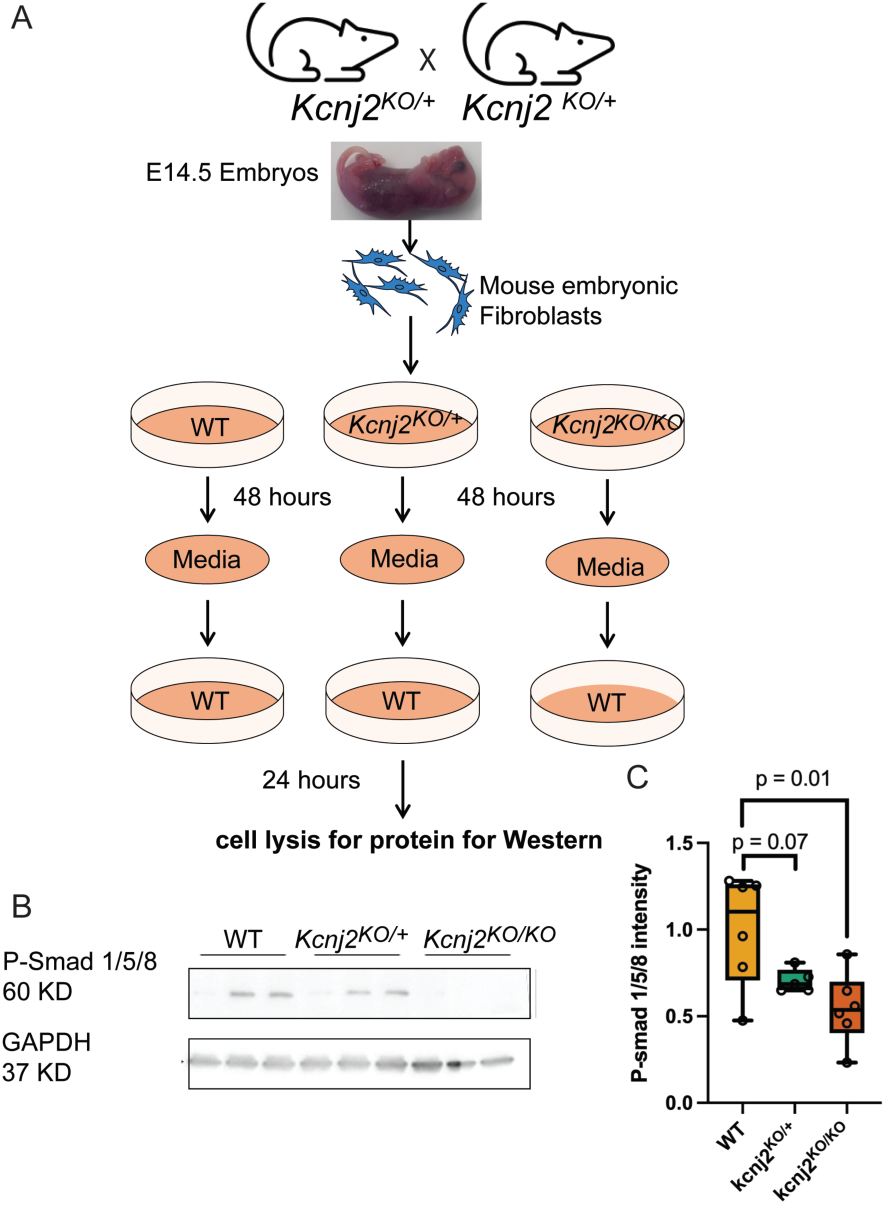
Deletion of a potassium channel that sets resting membrane potential reduces BMP in conditioned media. (A) A diagram shows the experimental design. Media was conditioned for 48 hours with MEFs from E14.5 WT, *Kcnj2^KO/+^*, or *Kcnj2^KO/KO^*. Conditioned media was placed on WT MEFs for 24 hours. Then MEFs were lysed, and protein was isolated for western blot analysis. (B) Representative western blot showing anti-rabbit P-Smad 1/5 and anti-GAPDH (loading control) in WT lysates incubated in conditioned media from MEFs from three different embryos of each genotype. (C) A graph shows quantification of relative fluorescence of P-Smad 1/5/8 from western blots (p values attained via T-Test, N=6 WT embryos, 5 *Kcnj2^KO/+^*, and 6 *Kcnj2^KO/KO,^*.

### Development of BMP4 release reporter

To examine the release of BMP4 from within BMP-producing mammalian cells, super ecliptic pHluorin (SEP), a pH-sensitive GFP variant, was inserted into the linker domain of BMP4 (Fig. 2A, see methods), similar to the imaging tool used previously in *D. melanogaster* (*28*). The fluorescence of SEP is quenched in acidic conditions, such as within a vesicle, and fluoresces in neutral conditions, such as when released into the extracellular environment. To test the function of this BMP4-SEP reporter, we utilized an immortalized mouse embryonic palatal mesenchyme (iMEPM) cell line (*45*). iMEPM cells are immortalized cells with an increased proliferative capability but maintain palatal mesenchyme characteristics such as morphology, transcriptional landscapes, and migratory capability (*45*). These cells are representative of the E13.5 mouse palate at an important time for signaling in the developing palate. iMEPMs can be used to investigate BMP signaling in mammals. To test the function of the reporter, iMEPM cells expressing BMP4-SEP were cultured for twenty-four hours. Ammonium chloride (5 mM) was added to the cells during live imaging to neutralize the cellular pH without depolarizing them (Fig. 2B), and the resulting changes in SEP fluorescence were recorded (Fig. 2C) (46). Upon neutralization, the iMEPM cells transfected with the BMP4-SEP construct had a robust increase in fluorescence, and this increase was significantly higher (p=0.0019) than that of our empty vector control plasmid (pCIG-GFP) (Fig. 2C). These results are consistent with our expectations of the reporter construct and thus confirm the validity of the tool.

**Figure 2:**
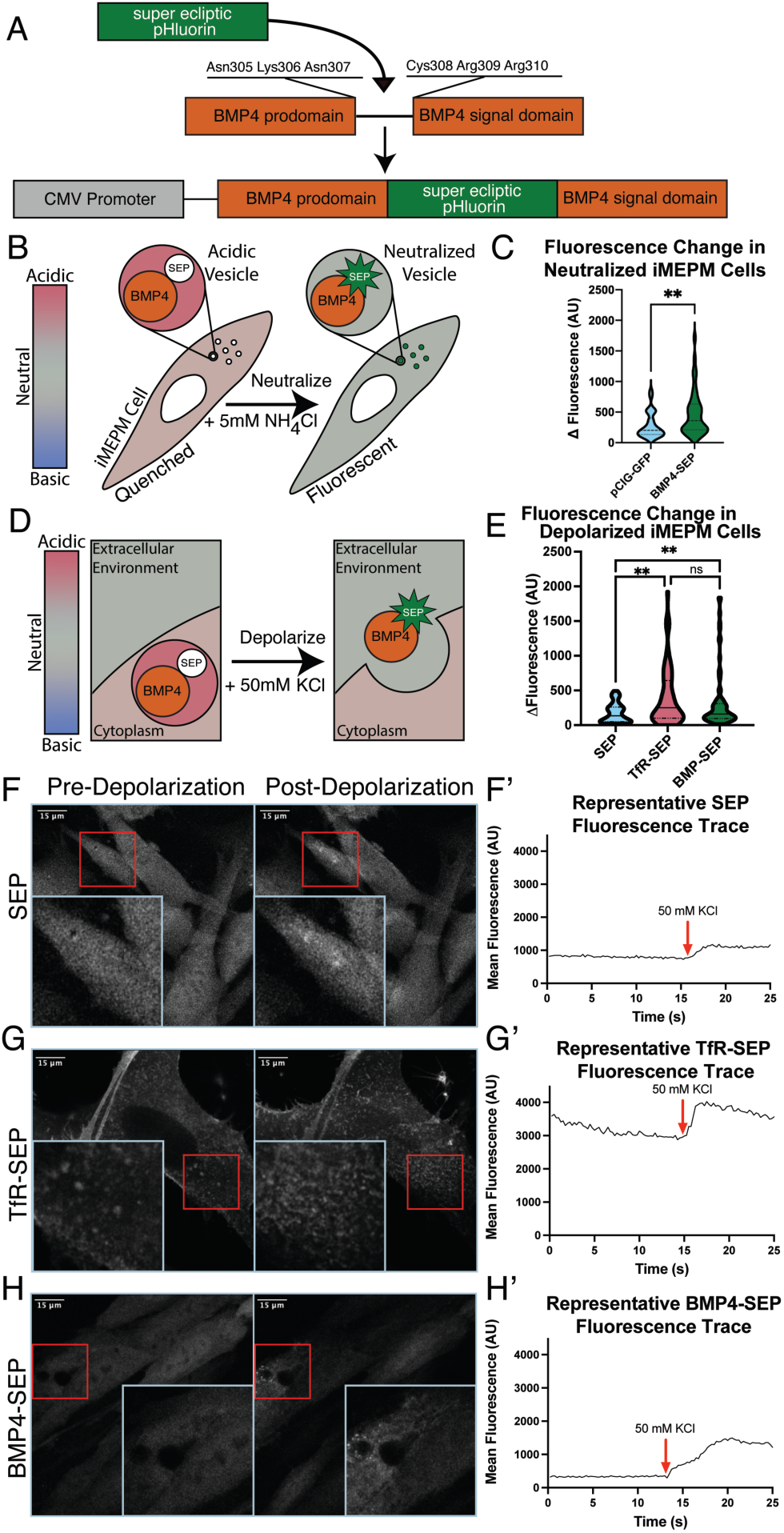
Depolarization of transfected iMEPM cells induces BMP4 release. **(A)** Diagram of BMP4-SEP fusion protein. Super ecliptic pHluorin (SEP) was inserted into the linker domain of BMP4 between Asn307 and Cys308 and cloned into a plasmid under control of a CMV promoter. **(B)** Diagram of iMEPM neutralization by ammonium chloride (NH_4_Cl). The acidic environment of a vesicle (red, pH ∼ 5.5) restricts the fluorescence of SEP. When NH_4_Cl is applied to a cell, the environment becomes neutralized (pH = 7.4), unquenching SEP and allowing for increased visualization of fluorescence. **(C)** A violin plot shows a significant increase in fluorescence amplitude between iMEPM cells expressing BMP4-SEP compared to a pCIG-GFP control after neutralization by the addition of 5mM NH_4_Cl. (**p<0.005 by t-test) **(D)** Diagram of BMP4-SEP release in response to cellular depolarization induced by the addition of KCl. **(E)** ViolinPlot showing quantification of SEP fluorescence in iMEPM cells transfected with pcDNA-SEP (blue), TfR-SEP (pink) or BMP4-SEP (green). Changes in fluorescence amplitude in regions of interest (ROIs) taken in live imaging videos (see methods) were averaged and compared. BMP4-SEP and TfR-SEP transfected cells had significantly higher changes in fluorescence than pcDNA-SEP (both p < 0.0001) but were not found to be different from one another. **(F, G, H)** Representative images show pcDNA-SEP, TFR-SEP, and BMP4-SEP fluorescence in iMEPM cells pre- and post-depolarization by 50mM KCl. Red boxes denote the location of the magnified blue insets within each respective image. **(F’, G’, H’)** Representative fluorescence traces from iMEPM cells transfected with pcDNA-SEP, TFR-SEP, or BMP4-SEP construct. The addition of 50mM KCl is denoted with the red arrow. (**p<0.002, ns=not significant by 2-way ANOVA)

### Depolarization induces vesicular release in iMEPMs

To test the hypothesis that palate mesenchyme cells are capable of depolarization-induced vesicular fusion for BMP release, we live imaged iMEPMs expressing a canonical marker of vesicular fusion, transferrin receptor-super ecliptic pHluorin (TfR-SEP) (*47, 48*). We compared iMEPMs expressing TfR-SEP and SEP alone as a control, during induced depolarization events (Fig. 2D). Fluorescence is dispersed throughout the cytoplasm of iMEPM cells that express SEP alone (Fig. 2F, Video S1). Upon depolarization, induced by addition of 50 mM potassium chloride, we did not see stark increases in fluorescence in cells expressing SEP alone (Fig. 2F, F’, Video S1). In contrast, depolarization of TfR-SEP expressing cells significantly increased punctate fluorescence intensity (Fig. 2G, G’, Video S2). This punctate increase in SEP fluorescence was significantly greater than iMEPM cells expressing SEP in the cytoplasm (176.6 ± 19.60 vs. 407.4 ± 54.71 AU, p=0.0034) (Fig. 2E). These data suggest that mouse palatal mesenchyme cells contain the machinery to respond to depolarization with vesicle fusion.

### Depolarization induces BMP4 release from iMEPMs

To determine if depolarization can induce BMP4 release, we depolarized BMP4-SEP expressing iMEPM cells during live imaging. Depolarization induced a clear increase in punctate BMP4-SEP fluorescence (Fig. 2H, H’, Video S3), supporting the model that depolarization can induce BMP4 release. Upon depolarization, BMP4-SEP fluorescence intensity increased significantly more than SEP in the cytoplasm in iMEPM control cells (176.6 ± 19.60 vs. 323.4 ± 46.42 AU, p=0.0016) (Fig. 2E). Interestingly, the amplitudes of fluorescence change for BMP4-SEP were not significantly different from those observed using the canonical TfR-SEP exocytosis reporter. Both BMP4-SEP and TfR-SEP fluorescence appeared as punctate on the surface of iMEPMs after depolarization (Fig. 2E, G-H’). These results suggest that depolarization induces BMP4 release from iMEPM cells.

### Depolarization increases BMP4 concentrations in iMEPM-conditioned media

An increase in BMP4-SEP fluorescence upon depolarization supports the hypothesis that depolarization can induce BMP release from iMEPM cells. To quantify and measure BMP release, we used a BMP ELISA. BMP4-SEP was transfected into iMEPM cells and conditioned media was collected before and after depolarization (Fig. 3A). Conditioned media collected after depolarization had BMP4 concentrations that were significantly greater than before depolarization (0.72 ± 0.08 vs. 0.95 ± 0.09 pg/mL, p=0.0003, N= 20 plates of cells) (Fig. 3B). The increase in BMP4 concentration in media following depolarization suggests that depolarization induces BMP4 release from iMEPM cells. These data suggest that ion channels that regulate the membrane potential of palate mesenchyme cells also control BMP release. We know that depolarization causes transient increases in cytoplasmic calcium to induce fusion of vesicles and mediate ligand release in excitable cells like neurons. We next asked if depolarization increases cytoplasmic calcium in the palatal mesenchyme.

**Figure 3:**
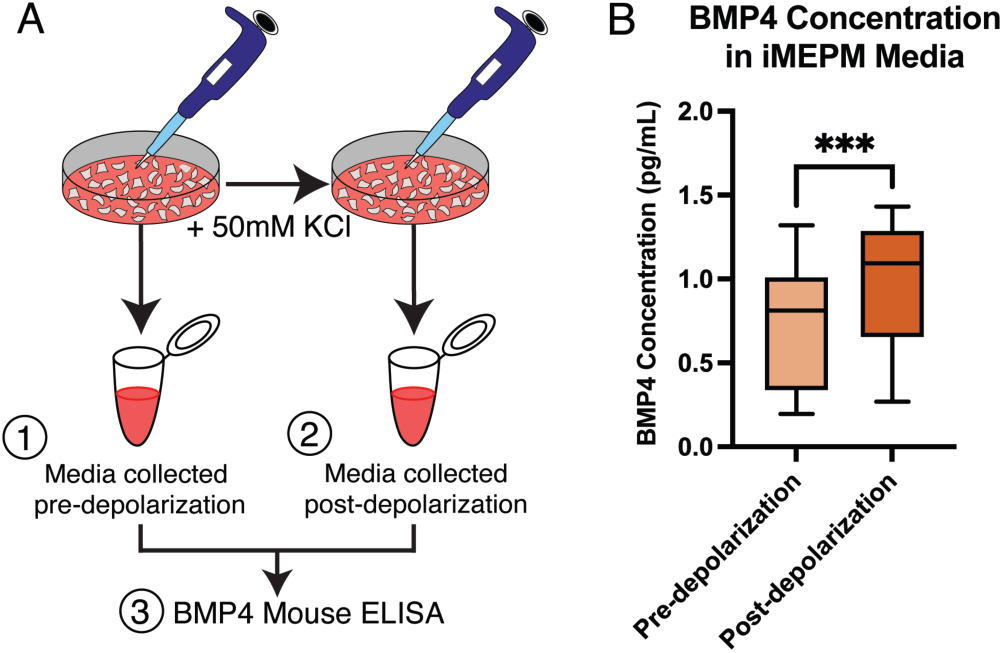
Depolarization increases BMP4 in conditioned iMEPM media. **(A)** A schematic shows the method used to quantify the amount of BMP4 in conditioned media before and after depolarization of BMP4-transfected iMEPM cells. iMEPM cells were cultured for 24 hours before conditioned media was collected and frozen. Cells were depolarized with KCl solution. Immediately following depolarization, conditioned media was collected and frozen. An ELISA was conducted with paired conditioned media samples collected before and after depolarization. **(B)** A paired box-and-whisker plot shows a significant increase in the BMP4 concentration after depolarization by KCl (***p<0.005 by T-test)

### Depolarization induces transient increases in cytoplasmic calcium in primary palatal mesenchyme

We expressed the fluorescent calcium reporter *GcAMP6s* in the palatal mesenchyme with the mouse *Wnt1Cre* driver. We dissected and cultured E13.5 palatal cells overnight. We imaged fluorescence over time with and without depolarization with KCl (50mM). Endogenous calcium events were observed in multiple fields of view within 3 independent cultured palates (Fig. 4A-D). Depolarization significantly increased GcAMP6s fluorescence (n=3 palates with 5-15 cells measured per palate, Fig. 4A-D, VideoS4). Importantly, depolarization did not decrease a cell’s ability to recover and exhibit additional calcium transients (Fig. 4C). We were also able to image and depolarize the same plate of cells multiple times when cells reacclimated in culture media for one hour between experiments. From this, we conclude that depolarization increases cytoplasmic calcium in E13.5 mouse palatal mesenchyme.

**Figure 4:**
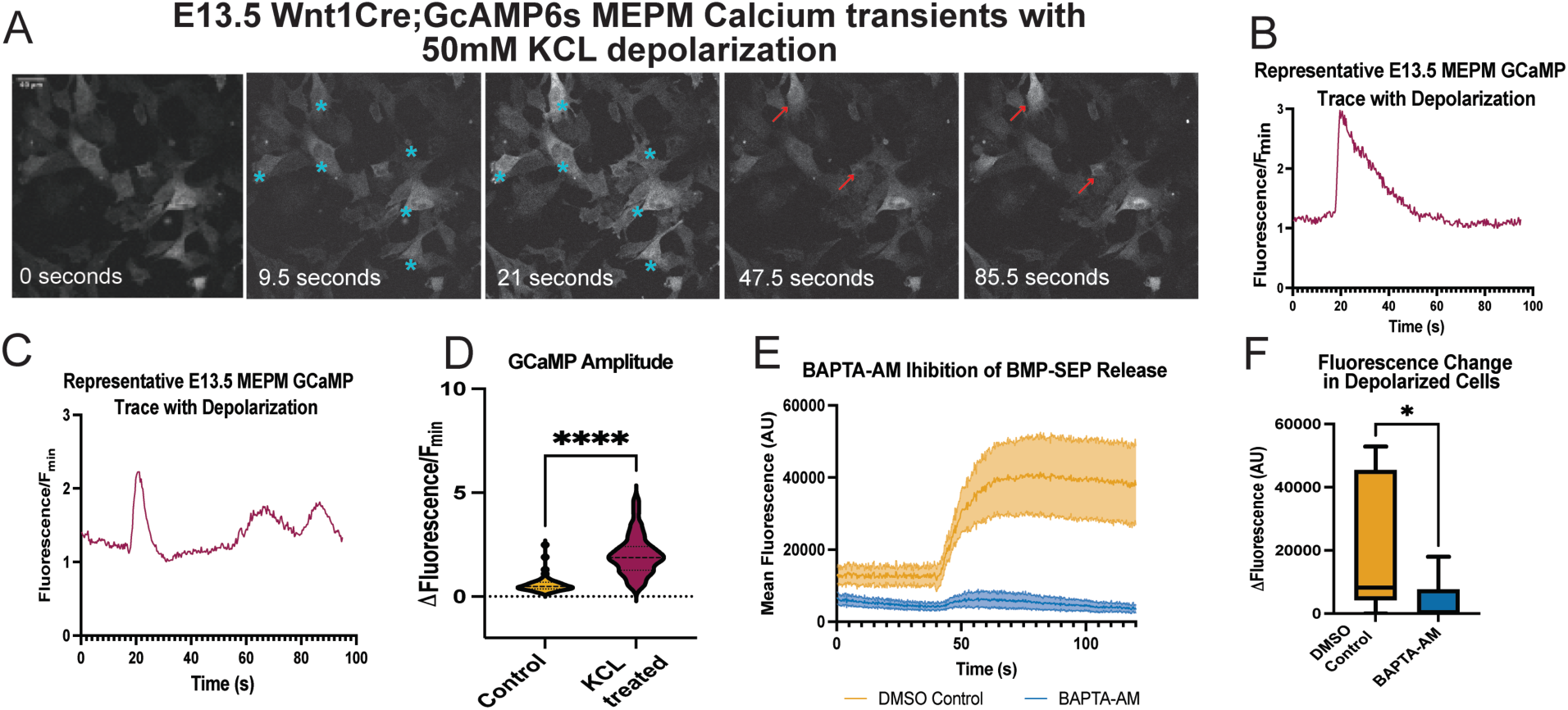
Depolarization induced BMP4-SEP release depends upon cytoplasmic calcium. **(A)** Representative images show that depolarization at 20 seconds increases the fluorescence of GCamp6 expressed in primary cultured E13.5 palate mesenchymal cells (blue stars), and cells have subsequent calcium release events following depolarization (red arrows). (**B**) Representative fluorescence profile of one cell over time. **(C)** Representative profile of fluorescence over time for a cell that undergoes a calcium transient with depolarization at 20 seconds followed by two endogenous transients. **(D)** Depolarization induces significant increases in fluorescence compared to background changes in fluorescence (****p<0.0005 by T-Test). **(E)** Mean fluorescence traces of BMP4-SEP release averaged between cells treated with or without BAPTA-AM. Yellow represents DMSO controls (n=6), and blue represents BAPTA-AM treated cells (n=10). The red arrow denotes the addition of isosmotic Tyrode’s media to induce depolarization. SEM is shown with shaded areas. **(F)** A box and whisker plot quantifying change in fluorescence amplitude between DMSO controls and BAPTA-AM treated iMEPM cells (*p<0.05 by T-test).

### Depolarization does not affect cell viability or future excitability

We measured cell viability with crystal violet assays after depolarization with potassium chloride. Cell viability was measured with crystal violet staining and showed intact, healthy iMEPMs after treatment with 50mM, 100mM, and 150mM KCl (Fig S1). Next, we asked if depolarization by our method disrupted native electrical activity of primary culture MEPMs. We show that depolarized cells have subsequent calcium release events (Fig 4C). Finally, after depolarization, we incubated cells in culture media overnight and imaged again the next day to confirm cells remained intact and excitable.

### BMP4 release from iMEPMs is calcium-dependent

To determine if depolarization-induced BMP release is dependent on cytoplasmic calcium, we repeated the BMP4-SEP experiments under isosmotic conditions with and without a cell-permeant calcium chelator called BAPTA-AM (Fig 4E-F). Cells were incubated in 100uM BAPTA-AM for one hour to chelate intracellular calcium (*49*). We found that incubation in BAPTA-AM reduced the number of cells that released BMP4-SEP upon depolarization from six out of twelve control cells to two out of ten treated cells. In the BAPTA-AM loaded cells, changes in the amplitude of BMP4-SEP fluorescence upon depolarization were significantly reduced from cells without BAPTA-AM treatment (19448 ± 6079 vs. 4092 ± 2141 AU, p=0.039) (Fig. 4E-F). Together, these data demonstrate that cytoplasmic calcium is necessary for depolarization-induced BMP release from palatal mesenchyme cells.

### Palate mesenchyme undergoes endogenous transient changes in intracellular calcium

For coordination of morphogenesis, cells need to send and receive precise signals. Neurons achieve precise temporal control of molecular signals using depolarization to control transient changes in intracellular calcium to drive vesicular fusion. To investigate whether transient changes in intracellular calcium occur at a time of active signaling for palatogenesis, we cultured E13.5 primary mouse embryonic palate (MEP) cells that express the calcium sensor, *GCaMP6s,* driven by *Wnt1Cre* and imaged fluorescence over several minutes. These experiments revealed calcium transients (Fig. 5A, Video S5) and demonstrated that calcium events in the palate can be periodic (Fig. 5B (yellow trace)). An average of 3.24 ± 0.16 events occurred in cells over the imaging period (384 seconds) (Fig. 5E). This number varied, however, with some cells experiencing only one event and others undergoing twelve events in that time. Calcium transients had an average fold change in amplitude (ΔFluorescence/F_0_) of 3.47 ± 0.091 ΔF/F_0_ (Fig. 5D). Calcium transients vary in intensity with some cells exhibiting extreme changes in GCaMP fluorescence while others show more mild changes (Fig. 5A, S2). In cells experiencing repeated periodic events, average inter-event period was 63.22 ± 2.50 seconds (Fig. 5F).

**Figure 5:**
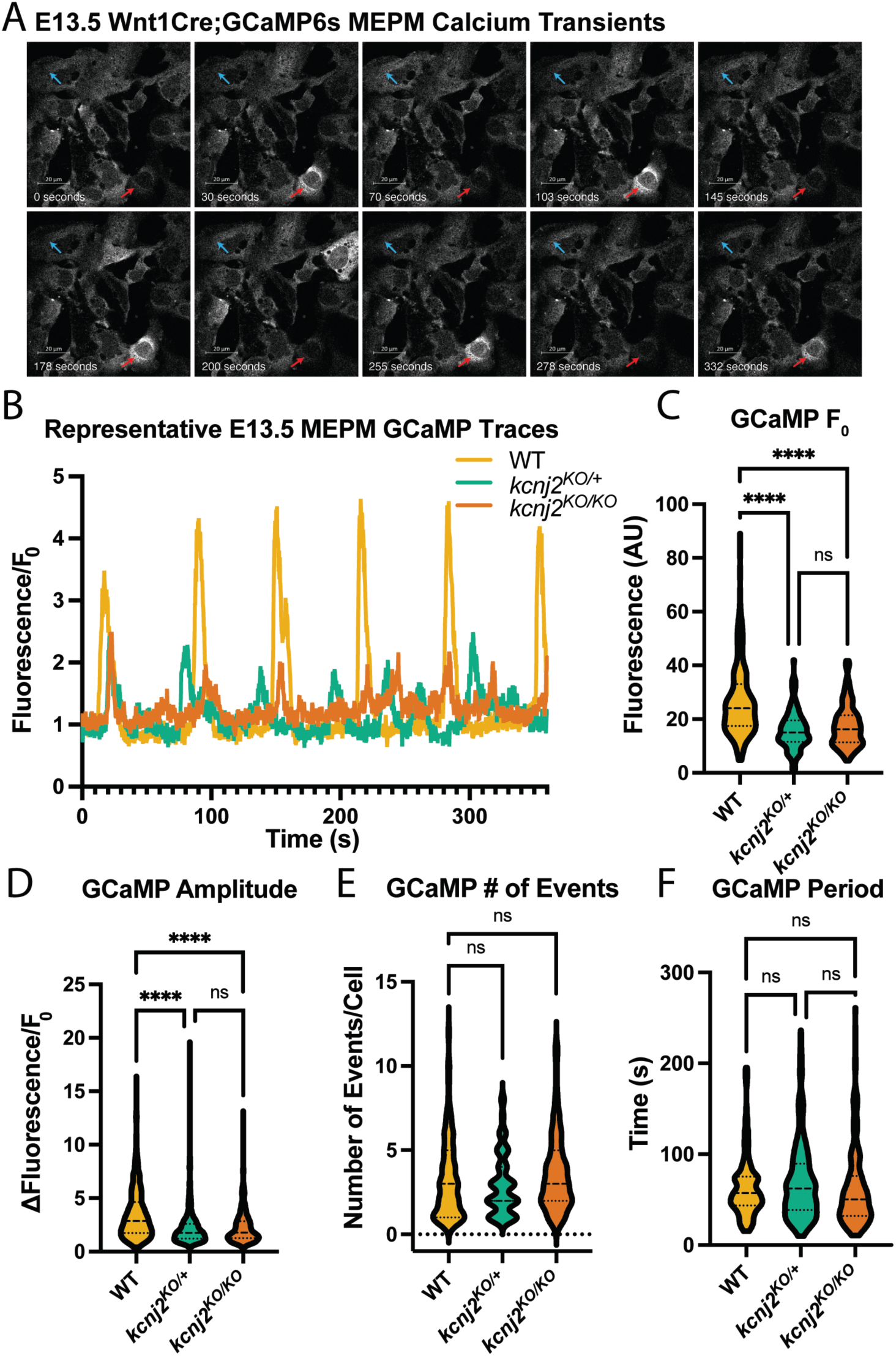
Calcium transients in E13.5 MEPM explants are regulated by *Kcnj2*. **(A)** *Wnt1CRE; GCaMP6s* expressing E13.5 MEP primary culture illustrating strong (red arrow) and weak (blue arrow) calcium activity in two cells. **(B)** Representative GCaMP6s fluorescence traces of *Wnt1CRE; GCaMP6s* (yellow), *Wnt1CRE; GCaMP6s; Kcnj2^KO/+^* (green), and *Wnt1CRE; GCaMP6s Kcnj2^KO/KO^* (orange) E13.5 MEPM cells. Quantification of GCaMP6s initial fluorescence **(C)** and amplitude **(D)** showing a significant reduction in *Wnt1CRE; GCaMP6s; Kcnj2^KO/+^*, and *Wnt1CRE; GCaMP6s Kcnj2^KO/KO^* compared to controls. **(E-F)** Quantification of GCaMP event period and number of events shows no change between controls. (****p<0.00001, ns=not significant by 2-way ANOVA)

### *Kcnj2* mediates endogenous calcium transients in mouse embryonic palate primary cell cultures

To determine if a channelopathy-associated ion channel (*Kcnj2*) regulates palatal calcium transients, we quantified calcium transients in primary MEPM cultures from *Wnt1Cre; GCaMP6s; Kcnj2^KO/+^* and *Wnt1Cre; GCaMP6s; Kcnj2^KO/KO^* mice compared to cells from *Wnt1Cre; GCaMP6s* control mice. When compared to *Wnt1Cre; GCaMP6s* littermates, loss of one or both copies of *Kcnj2* reduced the initial GCaMP6s fluorescence within the cells, defined as F_0_ (26.91 ± 0.97 vs. 15.82 ± 0.54 F_0_ in *Kcnj2^KO/+^*, p<0.0001, and 17.42 ± 0.61 F_0_ in *Kcnj2^KO/KO^*, p<0.0001) (Fig. 5C, Video S5-7). The mean amplitude of the calcium transients was similarly perturbed upon disruption of one or both copies of *Kcnj2* (3.47 ± 0.09 vs. 2.23 ± 0.08 ΔF/F_0_ in *Kcnj2^KO/+^*, p<0.0001, and 2.25 ± 0.07 ΔF/F_0_ in *Kcnj2^KO/KO^*, p<0.0001) palate cells (Fig. 5B, D). However, the interevent period of was not perturbed in either *Kcnj2^KO/+^* (63.22 ± 2.50 vs. 69.75 ± 2.78 seconds (s), p=0.63) or *Kcnj2^KO/KO^* (63.22 ± 2.50 vs. 62.55 ± 3.03 s, p=0.95) (Fig. 5F). Interestingly, the mean number of events per cell was not significantly different between conditions in either *Kcnj2^KO/+^* (3.24 ± 0.17 vs. 2.89 ± 0.14 events, p=0.63) or *Kcnj2^KO/KO^* (3.24 ± 0.17 vs. 3.60 ± 0.22 events, p=0.29) (Fig. 5E).

### Cranial neural crest cells undergo calcium transients that are controlled by Kcnj2

Because mutations in *Kcnj2* cause an array of craniofacial phenotypes in addition to cleft palate, we hypothesized that endogenous transient changes in intracellular calcium may be common to cranial neural crest (CNC) cells, the precursors of several craniofacial skeletal structures. To determine if transient changes in intracellular calcium occur in CNC cells, we cultured and live imaged E9.5 *Wnt1Cre, GCaMP6s* primary CNC cells. Live imaging of CNC cells that express GCamp6s revealed periodic calcium transients in single cells with an average amplitude of 1.80 ± 0.14 F/F0 (n=89 events) (Fig. 6A, C, D). Interestingly, we observed instances where groups of neighboring cells display synchronized increases in GCaMP6s fluorescence and in close succession. We call these synchronized or coordinated events calcium waves (Fig. 6B-C). To investigate if loss of *Kcnj2* perturbed CNC cells calcium transients, we compared GCaMP6s fluorescence in *Wnt1Cre; GCaMP6s* control mice to *Wnt1Cre; GCaMP6s; Kcnj2^KO/+^* and *Wnt1Cre; GCaMP6s; Kcnj2^KO/KO^*mice. We found that loss of both copies of *Kcnj2* significantly reduced the amplitude of GCaMP6s fluorescence (1.80 ± 0.14 vs. 0.91 ± 0.10 F/F0, p=0.019), Fig. 6D, while loss of only one copy of *Kcnj2* did not significantly decrease fluorescence amplitude (1.80 ± 0.14 vs. 1.60 ± 0.11 F/F0, p=0.93, Fig. 6D). Amplitudes of *Kcnj2^KO/KO^* CNC cells calcium transients were significantly lower than *Kcnj2^KO/+^* transients (p=0.04). We did not see a significant difference in F_0_ between any of the conditions (Fig. 6E). To determine if iMEPM cells undergo transient changes in intracellular calcium, we transfected iMEPM cells with a plasmid expressing *GCaMP6s*. We observed calcium events indicated by changes in GCamp6s fluorescence in iMEPM cells. (Fig. S3). These results support the hypothesis that CNC cells and their derivative cells undergo transient changes in intracellular calcium that may result in the release of BMP4.

**Figure 6:**
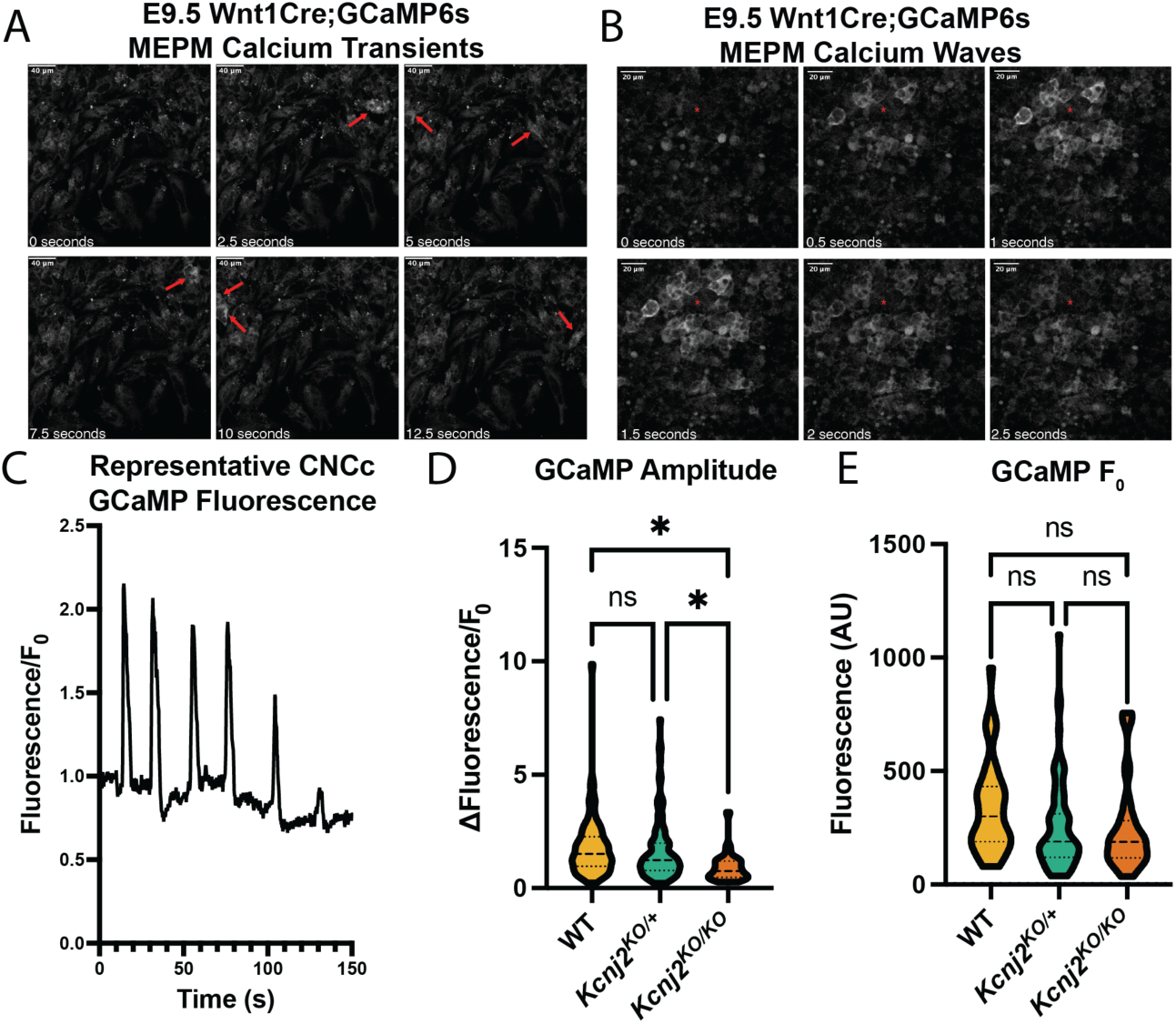
Disruption of *Kcnj2* in E9.5 MEPM explants. **(A)** *Wnt1CRE; GCaMP6s* expressing E9.5 CNC cells explants showing calcium transients (red arrows). **(B)** *Wnt1CRE; GCaMP6s* expressing E9.5 CNC cells explants showing a wave propagation event (centered on red asterisk). **(C)** An example of mean GCaMP fluorescence plot for an E9.5 CNC cells cell. **(D)** Quantification of GCaMP6s amplitude showing a significant reduction in *Wnt1CRE; GCaMP6s; Kcnj2^KO/KO^* compared to controls. **(E)** Quantification of GCaMP initial fluorescence shows no change from controls. (*p<0.0332, ns=not significant by 2-way ANOVA)

### Cranial neural crest-derived cells are electrically coupled

We observed that several cranial neural crest cells undergo calcium transients together, suggesting that they are electrically coupled. To quantify coupled electrical activity, we measured calcium activity throughout the area of the explant. We determined the percentage of the area that was synchronously electrically active. We found that 32% percent of the area underwent calcium transients together, suggesting that E9.5 cranial neural crest cells are electrically coupled (Fig. 7A-C). At later stages, palate mesenchyme cells are dissociated before they are cultured, but they may form connections again in cultured conditions. To determine whether cells at E13.5 are electrically coupled, we derived Pearson’s Correlation Coefficients for all possible cell pairs throughout the time course (Fig. 7E and H). Cell pairs with a correlation >0.5 were considered connected (Fig. 7D-I). The correlation between activity suggests an electrical coupling between pairs of neighboring cells (Fig. 7D-I). Analysis of individual time traces for regions showing the highest degree of coordinated activity indicates some synchronized activity even in far-apart cells (Fig. 7P). These data suggest that cranial neural crest cells and palate mesenchyme cells are electrically coupled.

**Figure 7:**
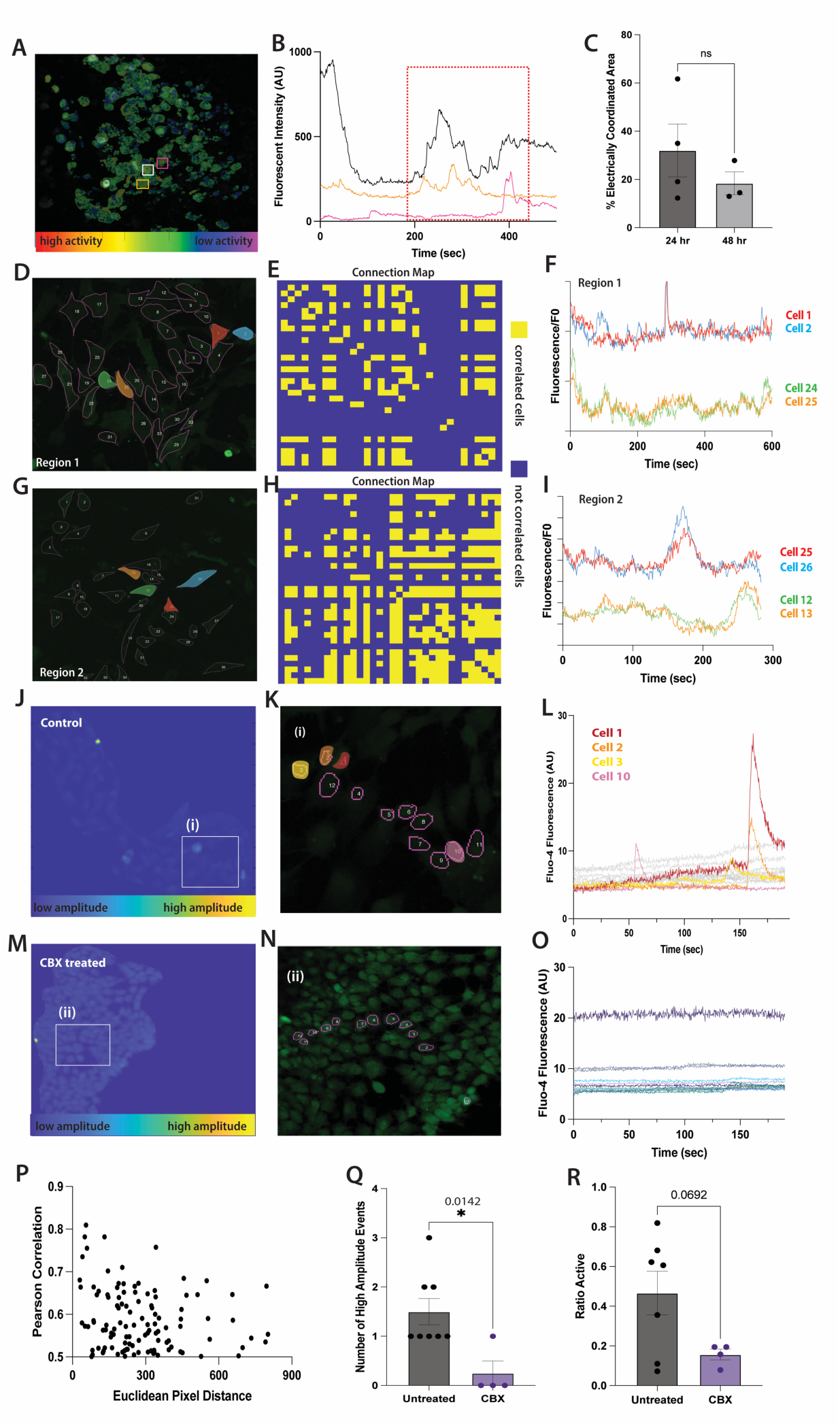
**(A)** A representative image shows cultured an E9.5 explant with three representative cells manually selected threshold of 0.5 **(B)** Normalized calcium timetrace of highly connected cells exhibiting a correlation coefficient > 60% of average connections across all cells **(C)** Percentage of area that shows highly correlated calcium activity within each field of view for 24 vs 48 hour samples (unpaired student t-test, p = 0.3630). **(D, G)** Cell maps of E13.5 palate mesenchyme cells manually selected ROIs from which calcium timetraces were extracted. **(E, H)** Highly connected cells exhibiting a correlation coefficient > 0.5 are indicated in yellow on Connection Maps**. (F, I)** Example calcium time traces exhibit instances of synchronized activity by highly connected cell pairs **(P)** Scatter plot comparing Euclidean pixel distance between cell nuclei and Pearson correlation coefficient showing a general decrease in correlation with increasing distance between cells (**J-K, M-N)** Confluent cells were hand selected as a region of interest from which the fluorescent calcium signal was extracted and analyzed on a pixel-by-pixel basis to obtain peak amplitude values among individual pixel time-courses. (**Q**). The number of high amplitude events was quantified for each sample. Untreated explants showed greater high amplitude events compared to CBX treated (unpaired student t-test, p = 0.0142) **(R)** Area active was determined as a proportion of pixel area that was both correlated and showed fluorescence above background signal. Explants treated with gap junction inhibitor CBX showed a decrease in area active compared to untreated (unpaired student t-test, p = 0.0692). **(L, O)** Individual calcium signal time courses are extracted for individual cells of regions showing the highest intensity peak amplitude. Individual cell activity was indeed greater in untreated samples compared to CBX treated as indicated by observed spike-like behavior shown in the signal.

### Gap junction inhibition reduces excitability

To determine if gap junctions contribute to the spreading of calcium between cranial neural crest cells, we measured correlated activity with and without gap junction inhibition. Carbenoxolone (CBX) is a widely used broad-spectrum gap junction antagonist. We used CBX to inhibit gap junctions in E9.5 explant cultures. CBX inhibition did not affect cell or explant viability, as shown by repeated experiments where CBX was washed off, explants cultured overnight, and imaged subsequent days. Explants were analyzed for high amplitude events and overall cell correlation to assess the effect of gap junction inhibition on calcium dynamics among explant tissue. We tested whether instances of high amplitude events decreased when gap junctions were inhibited via CBX (Fig. 7M-O). If gap junctions contribute to the production of increased calcium (indicated by high amplitude changes in fluorescence), then we expect high amplitude events to decrease with gap junction inhibition. Indeed, cells underwent significantly fewer high-amplitude events when treated with a gap junction inhibitor compared to untreated samples (Fig. 7L, O,Q). Next, we asked whether calcium correlation decreased when gap junctions were inhibited. Gap junction inhibition decreased correlated active area compared to control (Fig. 7R). When comparing clusters of cells including and surrounding regions of high amplitude, untreated explant cells showed more spiking or single oscillatory behavior compared to CBX treated (Fig. 7J-L). These results indicate a decrease in coordinated area and high amplitude events in confluent explant tissue treated with gap junction inhibitors (Fig. 7Q and R). These data support the possible role of gap junctions in explant tissue facilitating the accumulation of calcium and subsequent release of BMP.

### Ion channels are expressed throughout the palatal mesenchyme

To determine which ion channels are expressed in the palate mesenchyme and thus could regulate intracellular calcium at a time when BMP4 signaling is active, we identified ion channels that are expressed in an E13.5 anterior palate single-cell RNA sequencing (scRNA-seq) dataset produced in our lab (*50*). Mesenchymal and epithelial cell populations were identified by marker gene expression (*50*). We identified expression of ion channel encoding genes previously associated with craniofacial defects (*Cacna1a* and *Cacna1c*: CaV1.2, *Kcnj2*: Kir2.1, and *Gja1*: Cx43, Fig. 8). Excitingly, we detected expression of many other ion channels that could contribute to bioelectric signaling in the anterior palate mesenchyme (*Cacna1d*: CaV1.3, *Cacna1g*: CaV3.1, *Kcnb1*: Kv2.1, *Kcnb2*: Kv2.2, *Kcnc3*: Kv3.3, *Kcnn2*: SK2, Atp2a2: SERCA2, Stim1, Stim2, *Scn3a*: NaV1.3, *Scn8a*: NaV1.6, and *Gjc1*: Cxn45, Fig. 8). Interestingly, Kcnj2 is not highly expressed in the same palatal mesenchyme cells as BMP4 (Fig. 8R-T). However, virtually every cell in the anterior palate expresses at least one gene encoding gap junction proteins (Fig. 8U-W). Together with our observations that calcium transients are often coordinated between multiple cells, ubiquitous gap junction expression in the anterior palate suggests that these cells are electrically coupled and can propagate bioelectrical signals across long distances.

**Figure 8:**
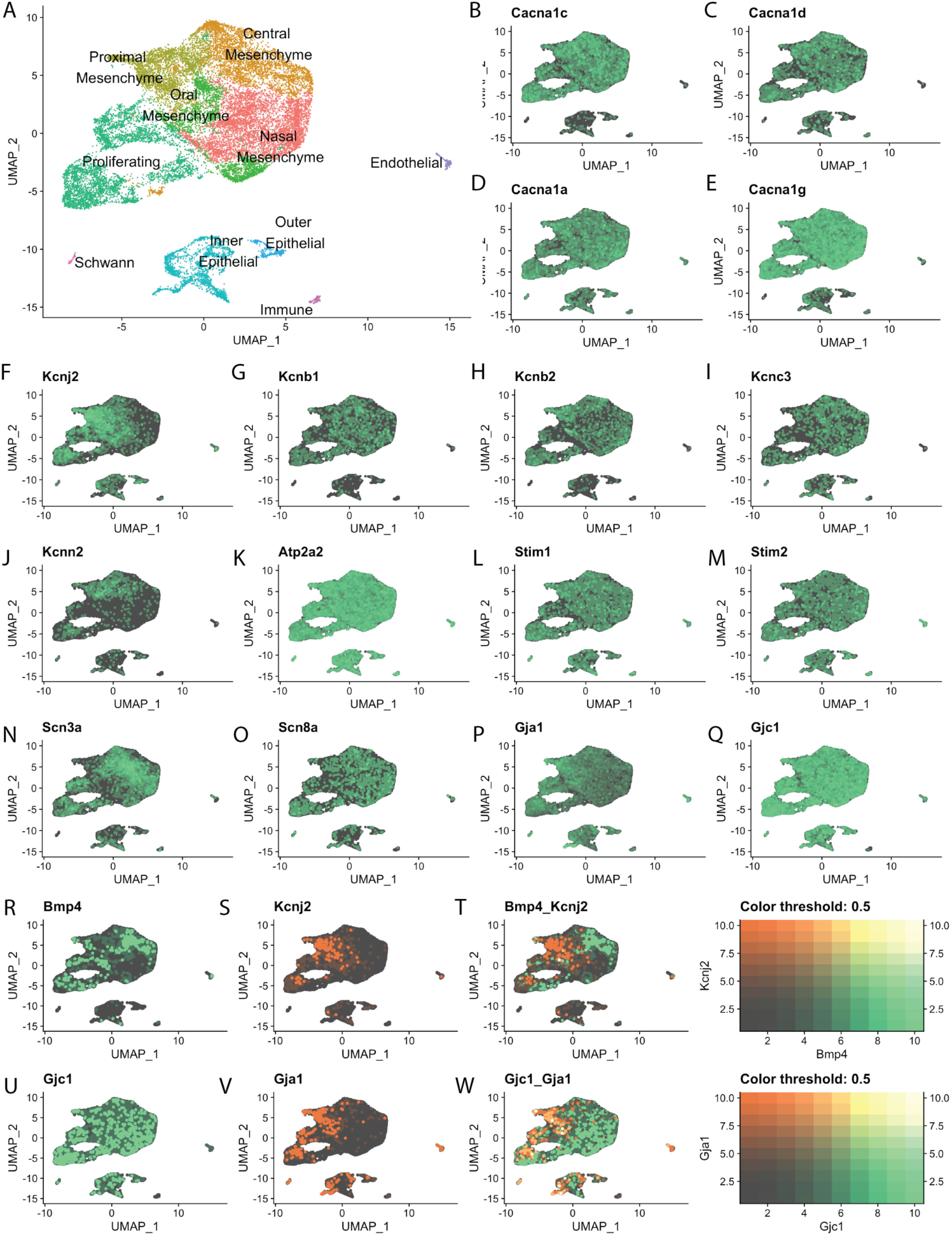
Ion channels and gap junctions are expressed in the E13.5 anterior palate. **(A)** UMAP detailing cluster identities adapted from Ozekin et al. FeaturePlots represent data from a single cell RNA sequencing of the E13.5 mouse anterior palate showing expression of ion channels and connexins in green with non-expressing cells in gray: **(B)** *Cacna1c* (Cav1.2, L-type calcium channel) **(C)** *Cacna1d* (Cav1.3, L-type calcium channel) **(D)** *Cacna1a* (Cav1.2, L-type calcium channel) **(E)** *Cacna1g* (T-type calcium channel) **(F)** Kcnj2 (Kir2.1, inwardly rectifying potassium channel) **(G)** *Kcnb1* (Kv2.1, voltage-gated potassium channel subfamily B) **(H)** *Kcnb2* (Kv2.2), **(I)** Kcnc3 (Kv3.3, voltage gated potassium channel subfamily C) **(J)** *Kcnn2* (KCa2.2, Potassium Calcium-activated channel subfamily N), **(K)** *ATP2a2* (SERCA2/ Atpase Sarcoplasmic/Endoplasmic Reticulum Ca2+ Transporting 2) **(L)***Stim1* (**M)** *Stim2* **(N)** *Scn3a* (Nav1.3, voltage gated sodium channel **(O)** *Scn8a* (Nav1.6 Voltage gated sodium channel) **(P)** *Gja1* (Gap junction protein alpha, Connexin 43) **(Q)** *Gjc1*(Gap Junction protein gamma 1, Connexin 45) **(R-T)** FeaturePlots of *Bmp4* (Bone morphogenetic protein 4) (green), *Kcnj2* (orange), and overlapped FeaturePlot of *Bmp4* and *Kcnj2*. Cells with high coexpression of both features will appear on a gradient to yellow. **(U-W)** FeaturePlots of *Gjc1* (green), *Gja1* (orange), and overlapped FeaturePlot of *Gjc1* and *Gja1*. Cells with high coexpression of both features will appear on a gradient to yellow.

## DISCUSSION

Here, we provide evidence that depolarization induces BMP release from mouse palatal mesenchyme cells. Depolarization-induced BMP release is dependent on cytoplasmic calcium. Furthermore, we show that palatal mesenchyme cells undergo transient changes in intracellular calcium, regulated by Kcnj2, a potassium channel required for palate development in mice and humans. This work suggests a mechanism linking observed craniofacial phenotypes of channelopathy patients with disruptions in traditional morphogen signaling. The necessity of ion channels for proper morphogenesis of human structures has been repeatedly documented (*KCNJ2*: Anderson-Tawil syndrome, *CACNA1C*: Timothy Syndrome, *GIRK2*: Keppen-Ludinski syndrome, *NALCN*: IHPRF1, and CHRNA7: 15q13.3 microdeletion syndrome) (*8, 9, 12, 13, 15, 30, 51–53*). Similarly, pharmacological and genetic disruption of ion channels during embryonic development causes morphological abnormalities in animal models (*54*). We propose that changes in membrane potential control secretion of vesicle-contained BMP ligands from palatal mesenchyme, in a calcium-dependent manner for craniofacial development.

Craniofacial phenotypes are not specific to disruption of one type of ion channel, suggesting that electrical activity mediated by several ion channels contributes to developmental signaling. Genetic and clinical evidence indicates that ion channels are important for BMP signaling (*27–29*). Mutations that disrupt *Kcnj2* cause craniofacial abnormalities such as dental defects, cleft lip and palate, hypertelorism, low-set ears, and micrognathia (small jaw) in human Andersen-Tawil syndrome (ATS) patients and mouse knockouts (*9, 13, 27, 29, 33, 55*). BMP signaling is required for the development of the structures affected in ATS patients. *Kcnj2* knockout (*Kcnj2^KO/KO^*) mice have similar craniofacial defects as ATS patients and as mice with disrupted BMP signaling (*27, 29, 43, 56–58*). Deletion of *Kcnj2* specifically in the cranial CNC causes similar craniofacial phenotypes as deletion of the BMP receptor - BMPR1a in the CNC (*56*). E13.5 *Kcnj2^KO/KO^* palatal shelves have decreased phosphorylation of Smad 1/5/8, a downstream target of BMP signaling and reduced BMP target gene expression (*27*). In the developing *Drosophila* wing disc, depolarization induces BMP/Dpp release (*28*). Our work suggests one possible conserved mechanism by which ion channels coordinate developmental signaling is by controlling release of BMP4.

We developed a novel tool to visualize BMP4 release from cells using the pH-sensitive GFP variant, superecliptic pHluorin (SEP). The BMP4-SEP release reporter is useful for the BMP research field enabling visualization of BMP release events. In some cells, we observe a slight increase in fluorescence in pcDNA-SEP expressing cells upon depolarization potentially due to small changes in cytoplasmic pH. In comparison, both the TFR-SEP and the BMP4-SEP underwent more profound increases in fluorescence in the presence of KCl. TFR-SEP is localized to vesicles and is commonly used as a marker of vesicular release, demonstrating that iMEPM cells, derived from the palate shelves of E13.5 mice (*45*), contain the proper machinery required for depolarization-induced vesicular fusion. Furthermore, we conclude that BMP4 is released from vesicles upon depolarization. The punctate appearance of TFR-SEP and BMP4-SEP fluorescence post-depolarization is consistent with vesicular fusion to the membrane. Future research is needed to determine the types of vesicles that contain BMP ligands and the specific mechanisms of release.

In excitable cells, such as neurons, if depolarization reaches a threshold at which voltage-gated Ca^2+^ channels open, increased Ca^2+^ levels change conformation of Soluble N-ethylmaleimide-Sensitive Factor Attachment Proteins REceptor (SNARE) proteins to induce vesicular fusion with the cellular membrane (*59*). If BMP-containing vesicle fusion is regulated by the same SNARE-dependent mechanism, sequestering intracellular Ca^2+^ would prevent an increase in BMP4-SEP fluorescence upon depolarization. A calcium chelator, BAPTA-AM, prevented any increase in BMP4-SEP fluorescence upon depolarization in most cultured palate mesenchyme cells. There were small increases in BMP4-SEP fluorescence upon depolarization in two BAPTA-AM treated cells which were significantly reduced in amplitude compared to cells that were not treated with BAPTA-AM. This suggests that BAPTA-AM treatment reduced the number of vesicle fusion events upon depolarization. Interestingly, in *Drosophila*, mutations in components of the SNARE protein complex have developmental defects (*60–62*). Perhaps there is a conserved role for calcium and SNAREs in morphogenesis.

Cells use chemo-transduction to instruct transcription in neighboring cells at a designated point in morphogenesis. In early murine development, Wingless-related integration sites (Wnts), Bone Morphogenetic Proteins (BMPs), Sonic hedgehog (Shhs), Fibroblast Growth Factors (FGFs) and other pathways initiate signaling cascades that ultimately affect transcription with cells to influence cell fate and ultimately morphogenesis (*63*). The temporal pattern of signaling affects the downstream transcriptional output in cell culture and in zebrafish osteoblast regeneration (*40*) (*41*). How do cells in developing tissues regulate these pathways to mediate the precise communication required for developmental decisions? Neurons, which also need precise communication, use depolarization as a critical step for vesicular release to regulate the release of molecular signals. We discovered cranial neural crest cells and palate mesenchyme cells, which are precursors to the palate, undergo rapid changes in intracellular calcium that are often cyclic and periodic, reminiscent of calcium spikes in neurons. Calcium transients are present in the neural crest precursor cells at E9.5, before BMP4 is expressed for palatogenesis. This raises the possibility that calcium transients may coordinate other signals in addition to BMP. Ion channel disruption affects other developmental pathways like Notch (*64–66*), Shh (*67*), and Wnt (*68–71*). Our work opens the question of whether electrical activity affects the extracellular presentation of these essential developmental signaling ligands as it does for BMP4.

We observed groups of cells undergo coordinated calcium transients suggesting that the cells are electrically coupled via gap junctions (Fig. 7). Electrical coupling would allow groups of cells to temporally coordinate periodic waves of BMP release across distances. Indeed, we found that gap junction genes are ubiquitously expressed within the E13.5 palatal mesenchyme (Fig. 8U-W). Expression of one of these gap junctions, *Gja1* (Cnx43), overlaps with expression of *Kcnj2* (Fig. 8S, U, W). Genetic disruption of Gja1 causes craniofacial phenotypes like that of *Kcnj2* perturbation (*72*). Gja1 modulates TGF-β1 and BMP2/4-mediated ERK signaling (*73*). ERK, a downstream target of BMP signaling, relies on periodic waves of activation to control osteoblast differentiation and regeneration of scales in Zebrafish (*41*). Our data show that inhibition of gap junctions in E9.5 palatal explants significantly reduces calcium activity in both active area and number of high amplitude calcium events (Fig 7Q and R). These results provide the basis for a mechanism explaining this observed temporal coordination of ERK signaling. In *Xenopus laevis* and *Danio rerio*, ion channels and gap junctions contribute to patterning and development suggesting that mechanisms controlling BMP ligand release could be further conserved (*26, 32, 74–76*). Furthermore, our data provides one potential explanation for why mutations in gap junction genes cause craniofacial defects in humans (*77–82*).

We hypothesized that ion channels that are important for craniofacial development regulate palatal intracellular calcium transients. Loss of one or both copies of *Kcnj2* caused an overall decrease in GCaMP fluorescence suggesting that basal calcium levels are reduced within these cells. Also, loss of Kcnj2 resulted in a significant decrease in calcium transient amplitude in palatal and CNC cells. Our data supports a model in which ion channels control membrane potential to mediate the secretion of BMP4. When ion channels are impaired, cells that rely upon those channels cannot regulate membrane potential, which disrupts intracellular calcium transients, propagation of these calcium signals, and consequently BMP secretion. Lack of regulated BMP secretion disrupts morphogenesis of the craniofacial complex (Fig. 9). This model explains a potential mechanism by which individuals with channelopathies have abnormal morphological development.

**Figure 9:**
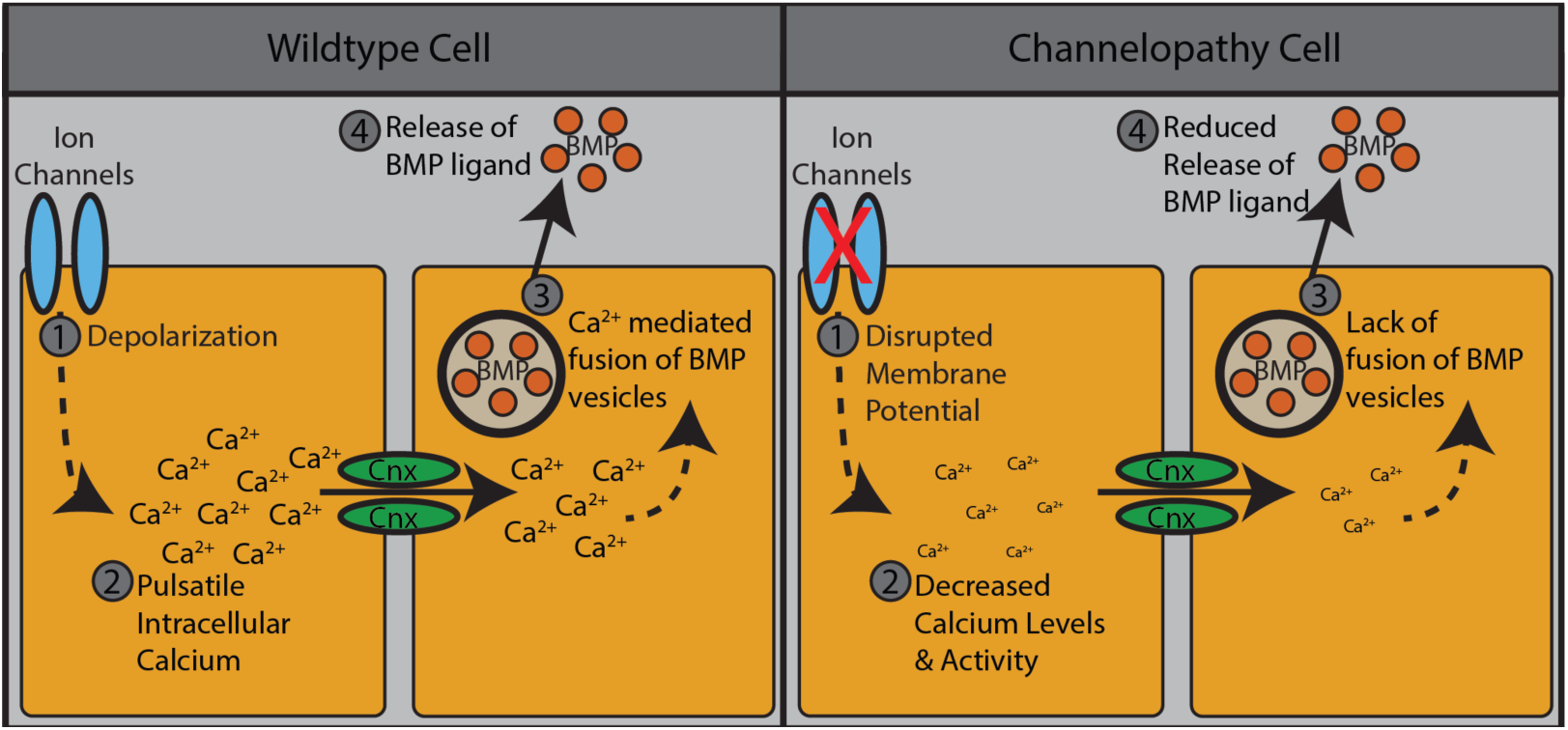
Proposed model of ion channel modulation of BMP secretion. In normal cells, ion channels regulate action potentials to control pulsatile intracellular calcium activity. This in turn drives fusion of BMP-containing vesicles allowing for release of BMP ligands. In channelopathy cells, action potentials are disrupted, resulting in decreased calcium activity and lack of fusion of BMP vesicles. This yields a reduced release of BMP ligands.

This research has clinical implications for fetal exposures to medications and recreational drugs that inhibit or activate ion channels. Fetal exposure to certain drugs that affect ion channel function increases the incidence of birth defects (*54*). For example, fetal exposure to an epilepsy medication and migraine medication called Topomax increases incidence of cleft palate (*85*). Topomax blocks voltage-dependent Na+ channels and AMPA glutamate receptors (*86–88*). Similarly, pentobarbital use during the first trimester increases incidence of cleft lip/palate in humans and in rodents and blocks GABA_A_ receptors, which are ligand-gated Cl^-^ ion channels (*19, 89*). In addition, Trimethadione, which blocks voltage-dependent T-type Ca^##^ channels, significantly increases the incidence of cleft palate, ear defects, and limb defects in the children of women who took the medication during the first trimester of pregnancy (*18, 90, 91*). We identify many previously uncharacterized ion channels expressed within the palatal mesenchyme that are targets of Pharmacologics (Fig. 8). Special care should be taken to understand how perturbation of these channels could affect palatal and craniofacial development. It may be that exposure to these drugs in utero disrupts cellular membrane potential to effect release of key morphogens, such as BMPs, thereby resulting in craniofacial defects.

## LIMITATIONS OF STUDY

While we show that BMP4 release can be induced by depolarization in cultured IMEPM cells and that primary cranial neural crest and palate mesenchyme cells undergo transient changes in intracellular calcium, we have not shown that depolarization induces BMP4 release in live mouse embryos. Furthermore, we cannot draw conclusions about whether BMP release is mediated by SNAREs as it is in neurons. Our work does not exclude the possibility that bioelectricity contributes to development via multiple mechanisms.

## ACKNOWLEDGEMENTS

We thank our colleague, Dr. Katherine Fantauzzo, for generously donating the iMEPM cells and Colleen Bartman for performing western blots. We thank Dr. Mark Dell’Acqua for insightful comments and suggestions.

## AUTHOR CONTRIBUTIONS

TI generated the BMP4-SEP plasmid. YHO and TI performed BMP4-SEP experiments. TI conducted iMEPM GCaMP experiments. YHO performed CNC and palate primary culture GCaMP experiments. MF performed and analyzed depolarization-induced increases in GCamp fluorescence in palate primary cultures. MF performed gap junction inhibition experiments in iMEPMs, palate primary cultures, and CNCs. CL analyzed waves of GCamp fluorescence and gap junction experiments. MF, TI, YHO, CL, and EAB created the figures and wrote first drafts of the components of the manuscript. EAB conceived and supervised experiments. EAB and YHO provided funding. All authors worked together in writing and editing the manuscript.

## SUPPLEMENTAL INFORMATION LEGENDS

**Supplemental Figure 1: Depolarization with KCl does not impair the viability of iMEPMs. (A)** A representative image shows untreated iMEPMs stained with Crystal Violet. (B) A representative image of iMEPMs treated with 10% hydrogen peroxide (known cell death inducer) shows that Crystal Violet staining is reduced when cells die and lose their adhesions. (C) A representative image shows that 100 mM KCl (the concentration we use for depolarization experiments) does not compromise iMEPM viability. (D) Absorbance quantification of Crystal Violet staining of iMEPMs with no treatment (media on iMEPMs), iMEPMs treated with hydrogen peroxide as a positive control, media with no cells, iMEPMs depolarized with 50 mM KCl, iMEPMs depolarized with 100 KCl, and iMEPMs depolarized with 150 KCl shows that depolarization with KCl does not compromise iMEPM viability.

**Supplemental Figure 2: GCaMP6s fluorescence traces of two MEPM cells from Figure 5A, color coded to denote cell, showing variability in GCaMP6s event amplitude.**

**Supplemental Figure 3: Representative GCaMP6s fluorescence trace from an iMEPM cell.**

**Video S1**: Cytoplasmic SEP fluorescence does not increase in a punctate pattern in depolarized IMEPM cells. IMEPM cells were transfected and cultured for 24 hours before imaging.

**Video S2**: TfR-SEP fluorescence increases in punctate pattern throughout iMEPM cells upon depolarization.

**Video S3**: BMP4-SEP fluorescence increases in a localized punctate pattern upon depolarization of iMEPM cells.

**Video S4:** GCaMP fluorescence increases with depolarization

**Video S5:** GCaMP fluorescence changes over time in primary cultures of wildtype mouse embryonic palate mesenchyme cells.

**Video S6:** GCaMP fluorescence changes over time in primary cultures of *Kcnj2^KO/+^* mouse embryonic palate mesenchyme cells.

**Video S7:** GCaMP fluorescence changes over time in primary cultures of *Kcnj2^KO/KO^* mouse embryonic palate mesenchyme cells.

## STAR METHODS

### BMP4-SEP cloning strategy

To generate a reporter of BMP4 release, we tagged BMP4 with superecliptic pHluorin (SEP), a pH sensitive GFP. To retain proper protein structure and function, SEP was inserted into the linker domain of BMP4 (RRKKNKN - SEP - CRRHSLYVDFSD), as previously described in *D. melanogaster* (Dahal, 2017). The BMP4-SEP construct was generated for our use by GenScript. The BMP4 feature of the plasmid pDONR233_BMP4_WT_V5, purchased from Addgene (Catalog #: 82937), was cloned into the pcDNA3.1(+) vector with a CMV promotor. A gBlock of SEP purchased from and generated by Twist Biosciences in San Francisco, California was sub-cloned into the linker domain described above. We confirmed the construct’s sequence using DNA sequencing through Quintara Biosciences.

### pcDNA3.1-SEP cloning strategy

A control plasmid for depolarization imaging was generated using SEP driven by a CMV promotor. The plasmid pcDNA3.1(+) (donated to us by a colleague) was digested with restriction enzymes HindIII and EcoRI to generate a linear plasmid. The plasmid TfR-mCherry-SEP (donated to us by Dr. Kennedy’s lab at the University of Colorado-AMC) was amplified via PCR using primers HindIII-SEP (CAGaagcttATGAGTAAAGGAGAAGAAC, purchased from IDT) and EcoRI-SEP (TCGgaattcTTATTTGTATAGTTCATCCA, purchased from IDT) to generate a SEP sequence with a HindIII overhang on the 5’ end and a EcoRI overhang on the 3’ end. The HindIII-SEP-EcoRI product and linearized pcDNA3.1(+) plasmid were then ligated together using the standard protocol from NEB T4 DNA Ligase (1:5 vector to insert ratio; purchased from NEB, catalog #: M0202S). Construct sequence was verified by DNA sequencing from Quintara Biosciences.

### iMEPM Cultures

iMEPM cells (immortalized mouse embryonic palatal mesenchyme) were cultured under the same conditions for each experiment. iMEPM cells were generated and donated by Dr. Fantauzzo’s lab at the University of Colorado, AMC and maintained by our lab. The media (referred to as iMEPM media from this point) used to grow the cells contained “DMEM with high glucose, no pyruvate or L-glutamine” (Gibco brand, purchased from ThermoFisher Catalog #: 11960044), 10% FBS (Gibco brand purchased from ThermoFisher catalog #: 16000044), 1% 200 mM L-Glutamine and 0.05% 50 mg/ul Pen-Strep. The cells were passaged several times post thawing and before splitting and plating for experiments. For depolarization imaging experiments, cells were cultured on WillCo Wells WillCo-dish® 35 mm glass bottom dishes (catalog #: HBST-3522) in 2mL of standard iMEPM media (@37C; 5% CO2). For the ELISA experiments, iMEPM cells were cultured in 35mm 6-well plates in standard iMEPM media (@37C; 5% CO2).

### MEF Isolation and Maintenance

Heterozygous *Kcnj2^ko/+^* mice were mated, and dams were considered E0.5 on the day a vaginal plug was observed. At E14.5, embryos were harvested. Heads were removed for phenotypic analysis. Internal organs were isolated for genotyping. Each embryo was dissociated with 3 mL of. 0.25% trypsin in EDTA. At 37. Degrees C for 5 minutes. Dissociated cells from each embryo were plated separately. We added 5 mL of a mixture of 10% DMEM, 1%FBS, 1% GlutaMAX (Gibco Life Technologies) and centrifuged for 5 minutes at 2000 g. The supernatant was replaced and resuspended in fresh 10% DMEM, 1%FBS, 1% GlutaMAX Pen Strep. Dissociated cells from each embryo were plated separately in 6-well plates.

### Western Blots

Protein isolated from WT cells treated with conditioned media was separated by gel electrophoresis and transferred to Turbo nitrocellulose transfer packs (BioRad). The blot was blocked for 1 hour in 5% milk/TBST shaking at room temperature. Following blocking, the blot was incubated in 1:1000 Rabbit Phospho-Smad 1/5/8 S463/465, (Cell Signaling #95165) in 5% milk/TBST shaking in 4°C overnight. The blot was incubated in 1:2000 secondary anti-rabbit for one hour at room temperature in the dark. The blot was washed four times in TBST for 15 minutes at room temperature before imaging with a BioRad Chemidoc MP Imaging System.

### Transfections

To examine the release of BMP4 from cells, we co-transfected iMEPM cells with either the BMP4-SEP construct and mCherry or pcDNA3.1-SEP and mCherry. Transfections followed the standard “Lipofectamine® LTX & PLUS™ Reagent” kit protocol (purchased from ThermoFisher, catalog #: 15338030). iMEPM cells were cultured in 6-well plates for ELISA experiments and on WillCo Wells WillCo-dish® 35 mm glass bottom dishes (catalog #: HBST-3522) for the depolarization imaging experiments. Cells were cultured until 80-90% confluent in standard iMEPM media, and media was then replaced with transfection solution. 1 mg of each type of DNA (BMP4-SEP and mCherry or PCDNA3.1-SEP and mCherry) was diluted in “Opti-MEM™ Reduced Serum Medium with no phenol red” (catalog #: 11058021) along with PLUS reagent and Lipofectamine-LTX from the transfection kit above. The solution was added to each plate dropwise and allowed to incubate (@37C; 5% CO2) for 4 hours, the solution was then replaced with iMEPM culture media and allowed to incubate overnight before imaging/ extraction (24 hours post transfection).

### Imaging and data analysis

Depolarization imaging was performed using a Zeiss LSM 880 confocal microscope with Airyscan. The images were taken as a 200 cycle time series at a speed of 250 ms with an average Airyscan scan time of 248.66 ms. To capture the SEP fluorescence, cells were excited at 488nm. To reduce file size while optimizing resolution, files were Airyscan processed after recording. Cells were imaged in iMEPM media. To capture the depolarization of the iMEPM cells, a solution of KCl (50 mM, based on the neuroscience field standard concentration for cell depolarization) was added to the dish via pipette at frame 50 while imaging. SEP fluorescence was measured before and after the depolarization event using Fiji, and amplitude was calculated for each event observed. Statistical analysis was done using the Student’s T-test function in GraphPad Prism8 to determine the significance of the difference in fluorescence.

### Isosmotic depolarization in BAPTA

Isosmotic depolarization and BAPTA-AM (ThermoFisher Scientific, #B1205) imaging was performed using a Zeiss LSM 900 confocal microscope with Airyscan II using the 40X water objective. Capture parameters were the same as the depolarization methods detailed above. To cause depolarization under isosmotic conditions, cells were imaged in a low potassium, high sodium Tyrode’s media (135 mM NaCl, 5 mM KCl, 2 mM CaCl_2_ · 2H_2_O, 1 mM MgCl_2_ · 6H_2_O, 25 mM HEPES, 10 mM Glucose, and 0.10% BSA, pH 7.4 with NaOH). At 150 frames, a high potassium, low sodium Tyrode’s media (5mM NaCl, 135 mM KCl, 2mM CaCl_2_ · 2H_2_O, 1 mM MgCl_2_ · 6H_2_O, 25 mM HEPES, 10mM Glucose, and 0.10% BSA, pH 7.4 with NaOH) was added directly to the plate to bring a the KCl to a final concentration of 50mM. Cells were exposed to either 100mM BAPTA-AM diluted in DMSO and Tyrode’s media or DMSO in Tyrode’s media alone for one hour at 37**°**C prior to imaging.

### ELISA and analysis

A BMP4-Mouse ELISA assay was performed to chemically confirm the presence of BMP4 in the ell media before and after depolarization. Cells were cultured and transfected with BMP4-SEP and mCherry as described above were depolarized with KCl (50mM). Fractions of media were collected before and after depolarization was induced. Fractions were prepared and assayed according to the standard protocol from the Abnova “BMP4 (Mouse) ELISA Kit” (catalog #: KA5051). The ELISA plate was read on the Biotek synergy H1microplate reader at 561 nm. Statistical Analysis was performed using the Students’ T-test function in GraphPad Prism8 to determine the significance of the difference in fluorescence intensity before and after depolarization.

### Crystal Violet assay and analysis

We followed the Cold Springs Harbor 2016 “Crystal violet assay for determining viability of cultured cells” protocol (M. Feoktistova, P. Geserick, and M. Leverkus). Briefly, iMEPM cells were seeded onto 96well plates, leaving at least three wells with no cells (media only). Cells were cultured overnight. Each KCl concentration was added to treatment wells in triplicate, leaving three seeded cells untreated as a negative control. Three wells were also treated with H2O2 to induce cell death as a positive control. KCl treatment was applied for 15 minutes to simulate experimental conditions of depolarization assays. The wells were then washed and stained with 0.5% crystal violet solution. Prior to imaging, the wells were washed and air-dried overnight. Optical density was measured at 570nm. Optical density values were graphed and used to calculate the percentage of light transmitted through the sample.

### GCaMP Methods

*Rosa26-GCaMP6s* (Jackson Laboratory, *B6;129S6-Gt (ROSA)26Sor^tm96(CAG-GCaMP6s)Hze^/J*) female mice were mated overnight with *Wnt1-CRE* males. Observation of a vaginal plug was considered day 0.5. On day 13.5, pregnant females were euthanized by isoflurane exposure and cervical dislocation. Embryos were dissected in ice-cold phosphate buffered saline (PBS) and limbs were removed for genotyping.

### Cell culture

Mouse embryonic palate (MEP) cells were collected from E13.5 embryos as per protocol adapted from [Bush and Soriano, 2010]. Briefly, paired palatal shelves were isolated from each embryo and placed in 100uL 0.25% trypsin/EDTA (ThermoFisher Scientific, #25200056) for 15 minutes at room temperature with frequent agitation. Trypsinization was stopped by adding 10 volumes of media (DMEM + 10% FBS). Glass bottom culture dishes (Willco Wells, HBST-3522) were prepared by coating in 0.005% fibronectin in PBS for 45 minutes at room temperature immediately before dissection. MEP cells were plated and cultured overnight at 37°C. The following day, cells were rinsed with PBS and fresh media was added.

### Calcium Imaging and Quantification

E9.5 CNC cells were imaged 24 hours post dissection on a ZEISS LSM 880 with airyscan confocal microscope using the 488 nm laser at 20X magnification at 4 Hz. over 2.5 minutes. E13.5 palate primary cultured cells were imaged 24 hours post dissection on a ZEISS LSM 900. GCaMP6s signal was observed using a 488 nm laser under 40X magnification at 3.16 Hz. All cells that showed high variance were selected as regions of interest (ROIs). Mean fluorescence profiles were plotted for each ROI over 6 minutes. Mean fluorescence profiles were used to obtain event numbers, event amplitude, and interevent period. An event was defined as a fluorescent spike two or more standard deviations above the mean fluorescence. Interevent periods were determined by measuring the time between the peaks of two adjacent amplitudes. Event numbers were calculated by counting the number of events, as defined above, within a cell. For each sample, a minimum of 5 fields of view (FOVs) were imaged.

### Quantification of GCaMP number of events, F_0_, amplitude, and interevent interval

Number of events were counted in each cell where activity was observed within a field of view (WT: N=3, n=198; Kcnj2^KO/+^: N=4, n=159; Kcnj2^KO/KO^: N=4, n=103). An event was defined as having a max fluorescence of two standard deviations or greater above mean fluorescence or the trace. Amplitudes were defined as the max peak fluorescence value of an event minus the minimum value before that peak (WT: N=3, n=633; Kcnj2^KO/+^: N=4, n=460; Kcnj2^KO/KO^: N=4, n=556). F_0_, or initial fluorescence, was determined by finding the minimum fluorescence value of the line trace (WT: N=3, n=199; Kcnj2^KO/+^: N=4, n=159; Kcnj2^KO/KO^: N=4, n=156). The interval between two events within the same cell was measured as the interevent period (WT: N=3, n=172; Kcnj2^KO/+^: N=4, n=237; Kcnj2^KO/KO^: N=4, n=208). For iMEPM experiments, the average change in fluorescence was not quantified due to differences in GCamp plasmid copy number.

#### Gap Junction inhibition calcium signal analysis (Figure 7A-C, M-P)

Explants were imaged at 350 ms/frame and analyzed using MATLAB. Confluent cells were hand selected and analyzed. Peak amplitude of fluorescence was identified for each pixel and plotted as a heatmap relative to the mean where yellow indicates high amplitude and blue corresponds with low. Number of high amplitude events was quantified based on regions within the heatmap showing high peak amplitude. Average correlation was assessed on a pixel-by-pixel basis and superimposed with the peak amplitude map to obtain ratio of ‘active’ area with respect to entire tissue area. Specific cell clusters were hand selected and assessed based on areas of high peak amplitude whereby calcium signal was extracted using MATLAB and plotted to compare against CBX treated samples.

### Network analysis

Calcium was analyzed using MATLAB scripts designed to identify individual cells with highly correlated activity (**Figure 7D-L**). First, single cells were identified by hand. Individual calcium time traces were extracted and normalized. Pearson’s correlation coefficient was derived for all identifiedcell pairs. Pairs with a correlation >0.5 were considered “connected” and mapped as a Boolean matrix.The population of cells per link were categorized and ordered as a percentage of total connections. The top 60% of cells were considered highly correlated.

### scRNA-Sequencing Analysis

UMAP and FeaturePlots were generated using Seurat on a publicly available dataset profiling the E13.5 anterior palatal mesenchyme (*54*).

