## Supplementary figures and images for "Depolarization induces calcium-dependent BMP4 release from mouse embryonic palate mesenchyme"

### Supplemental Figure 2

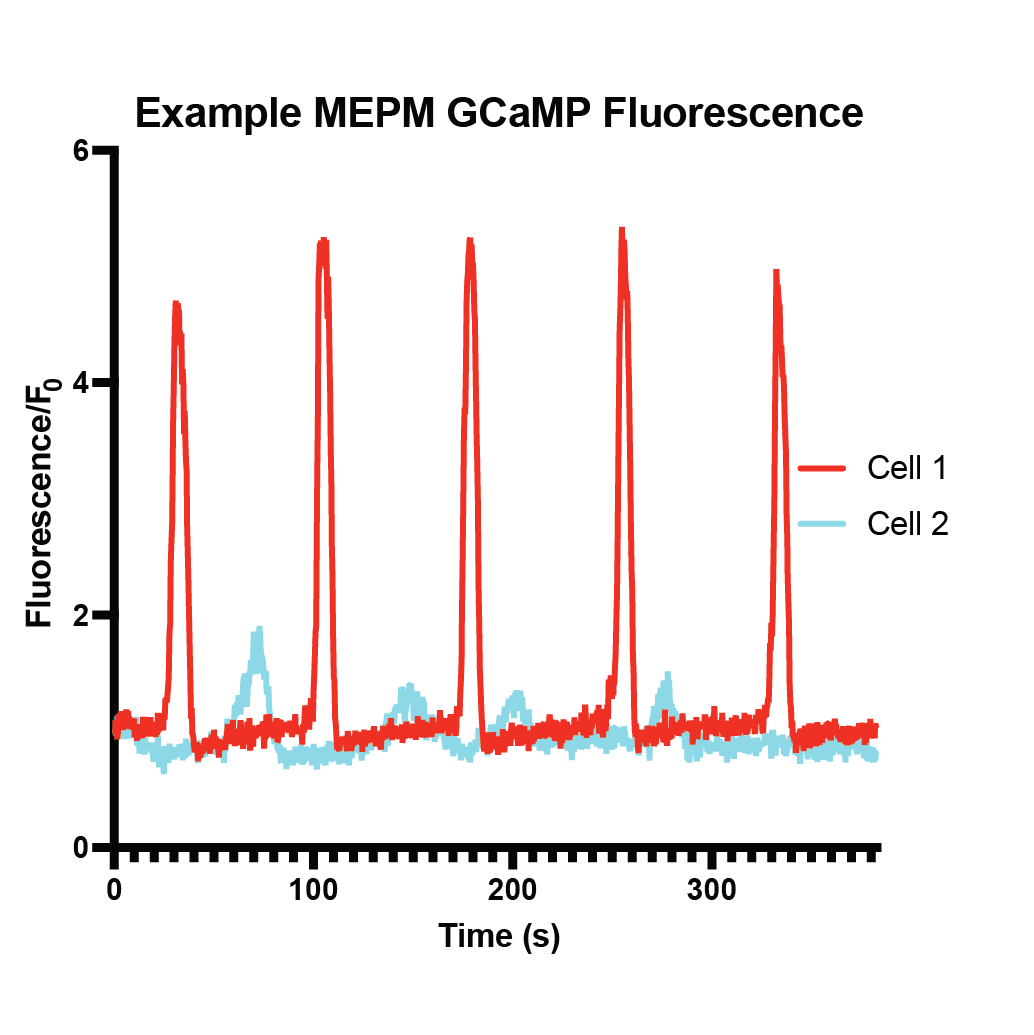

### Supplemental Figure 3

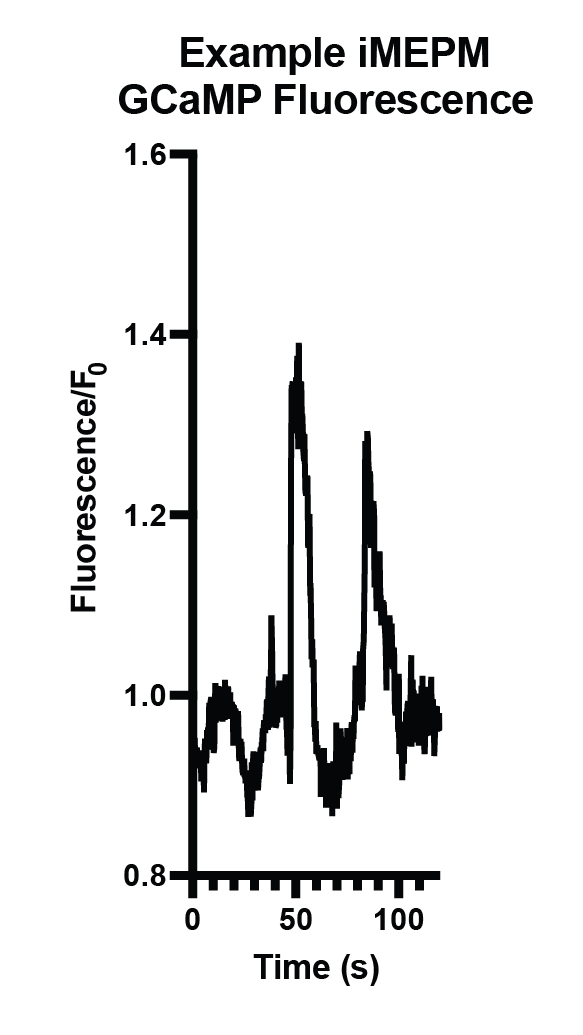
